# Myelin imaging of the basal forebrain in first-episode psychosis

**DOI:** 10.1101/2021.09.12.459966

**Authors:** Min Tae M. Park, Peter Jeon, Leon French, Kara Dempster, M. Mallar Chakravarty, Michael MacKinley, Julie Richard, Ali R. Khan, Jean Théberge, Lena Palaniyappan

**Affiliations:** Department of Psychiatry, Schulich School of Medicine and Dentistry, Western University, London, Canada; Department of Medical Biophysics, Western University, London Canada; Robarts Research Institute, Western University, London, Canada; Lawson Health Research Institute, London, Canada; Department of Psychiatry, University of Toronto, Toronto, Canada; Department of Psychiatry, Dalhousie University, Halifax, Canada; Departments of Psychiatry and Biological and Biomedical Engineering, McGill University, Montreal, Canada; Cerebral Imaging Centre, Douglas Mental Health University Institute, Montreal, Canada

## Abstract

Cholinergic dysfunction has been implicated in the pathophysiology of psychiatric disorders such as schizophrenia, depression, and bipolar disorder. The basal forebrain (BF) cholinergic nuclei, defined as cholinergic cell groups Ch1-3 and Ch4 (Nucleus Basalis of Meynert; NBM), provide extensive cholinergic projections to the rest of the brain. Here, we examined microstructural neuroimaging measures of the cholinergic nuclei in patients with untreated psychosis (∼ 31 weeks of psychosis, <2 defined daily dose of antipsychotics) and used Magnetic Resonance Spectroscopy (1H-MRS) and transcriptomic data to support our findings. We used a cytoarchitectonic atlas of the BF to map the nuclei and obtained measures of myelin (quantitative T1, or qT1 as myelin surrogate) and microstructure (axial diffusion; AxD). In a clinical sample (n=85; 29 healthy controls, 56 first-episode psychosis), we found significant correlations between qT1 and 1H-MRS-based dorsal anterior cingulate choline in healthy controls, while this relationship was disrupted in FEP. Case-control differences in qT1 and AxD were observed in the Ch1-3, with increased qT1 (reflecting reduced myelin content) and AxD (reflecting reduced axonal integrity). We found clinical correlates between left NBM qT1 with manic symptom severity, and AxD with negative symptom burden in FEP. Intracortical and subcortical myelin maps were derived and correlated with BF myelin. BF-cortical and BF-subcortical myelin correlations demonstrate known projection patterns from the BF. Using data from the Allen Human Brain Atlas, cholinergic nuclei showed significant enrichment for schizophrenia and depression-related genes. Cell-type specific enrichment indicated enrichment for cholinergic neuron markers as expected. Further relating the neuroimaging correlations to transcriptomics demonstrated links with cholinergic receptor genes and cell type markers of oligodendrocytes and cholinergic neurons, providing biological validity to the measures. These results provide genetic, neuroimaging, and clinical evidence for cholinergic dysfunction in schizophrenia and other psychiatric disorders such as depression.

## Introduction

Dysregulation of the cholinergic system has been long suspected in the pathophysiology of psychotic disorders such as schizophrenia(1, 2). Several lines of evidence support a cholinergic imbalance in psychosis(3), particularly in the negative symptoms such as psychomotor retardation and inattention(4, 5). The cholinergic hyperactivity induced by physostigmine worsens negative symptoms(4), while anticholinergics provide some relief for negative symptoms(6, 7). Muscarinic (M1/M4) agonist xanomeline has been shown to have clinical efficacy in schizophrenia(8, 9). In line with these findings, post-mortem human studies have identified a biological basis for these observations; specifically, the distributed reduction of muscarinic receptors in a subset of patients(10, 11). Taken together, these finding motivate calls for focused investigations of the cholinergic system to aid therapeutic discoveries in psychosis(9).

Despite the substantial evidence for a cholinergic abnormality in schizophrenia, it is not clear how a disrupted cholinergic system relates to the first presentation of psychosis, before treatments with anticholinergic effects are started. The three major hurdles in this regard are (1) the challenges in non-invasive study of the basal forebrain (BF), a structure that provides cholinergic projections to extensive areas of the cortical mantle; (2) the difficulties in direct quantification of acetylcholine(12) through in vivo imaging studies, (3) challenges in studying symptomatic untreated patients without the confounds of illness chronicity and long term antipsychotic exposure(13).

While there are a number of whole brain morphometric studies(14), and a region-of-interest study(15) of BF in schizophrenia, none have identified notable changes in the basal forebrain. A recent study specifically assessed gray matter volume of the BF in schizophrenia, while results were not significant after taking into account global effects(16). This contrasts with Alzheimer’s disease, where BF volume reduction is seen preceding other degenerative changes across the brain(17-19) suggesting general vulnerability of the region in neuropsychiatric conditions. This vulnerability of BF has been linked to its later onset of myelination and the relative sparsity of myelin sheaths in the BF compared to other regions of the brain(20-22). Higher myelin concentration may reflect reduced metabolic demands on the ensheathed neurons, and thus higher resilience to degenerative processes(23).

In a focused post-mortem examination of Nucleus Basalis of Meynert (NBM), Williams and colleagues(24) identified a notable reduction in oligodendrocyte density and glial cell abnormalities in schizophrenia. It is unknown if these changes are restricted to the NBM, which primarily projects to the neocortex, or if they extend to anteromedial BF nuclei (Ch1-3, or Broca’s diagonal band) that project to the hippocampus(25). While the relative lack of grey-white matter differentiation in the basal forebrain often limits accurate volumetric measurements, advances in human neuroimaging have provided us with probabilistic maps of BF based on cytoarchitectonic studies(26). Though oligodendrocytes cannot be directly measured using MRI in humans, a number of related microstructural properties can be examined in vivo. Of particular relevance is quantitative T1 or qT1 which has a high negative correlation with myelinated axon fraction, while axial diffusivity (AxD) best captures the non-myelinated axon fraction; both qT1 and AxD increase in experimental animal models of demyelination(27).

Acetylcholine and free choline, the precursor of acetylcholine, together contribute to a small portion of the magnetic resonance spectroscopy (MRS) total choline spectra, but this signal is not separable from that of other membrane bound choline moieties(28). Nevertheless, several observations suggest that the variations in the MRS choline signal may reflect variations in cholinergic tone. Xanomeline, a muscarinic agonist, decreases MRS choline resonance in Alzheimer’s Disease patients(29). In mice, the anticholinergic scopolamine induces an acute reduction in MRS choline signal that returns to baseline in 72 hours(30), while anticholinesterase donepezil increases choline resonance(31). In rats, MRS choline signal intensity shows a high degree of correlation with direct tissue measurement of acetylcholine levels across various brain regions(32). In humans, free choline, when teased apart from bound choline, changes with the performance of tasks such as reversal learning that are dependent upon cholinergic transmission(28, 33). While these fMRS studies have focussed on striatal choline signal, the diffuse prefrontal cholinergic projections from the basal forebrain(34-36), indicate that the overall cholinergic tone of BF projection are best estimated from the frontal cortical regions. MRS choline is increasingly being used as a proxy measure for acetylcholine levels in the anterior cingulate cortex in patients with psychosis(37).

The anterior cingulate cortex (ACC) is an important site of cholinergic projection from the basal forebrain in animals(38) and humans(25, 39). In particular, the NBM is specifically connected to the dorsal ACC component of the salience network(25), enabling contextual integration and cognitive control function(40). The Salience Network, in turn, plays a critical role in the resource allocation for stimulus evaluation and action outcomes that involve the deployment of large-scale cortical networks(41). Prior research, from our groups and others(42-45), has implicated the SN in the pathophysiology of psychosis. Given the cholinergic hypothesis of psychosis, and the SN dysfunction observed in this illness, it is likely that the structural integrity of basal forebrain cholinergic nuclei influences the cholinergic tone of the SN, but this has not been evaluated to date.

Here, we examine the microstructure of the basal forebrain cholinergic nuclei in first-episode psychosis (FEP) using ultra-high imaging. We quantified choline resonance from the dorsal ACC using 7T proton MRS. We related this resonance to BF qT1 in 56 patients with first episode psychosis and 29 healthy individuals, anticipating a dissociation between the microstructure of BF and dorsal ACC choline levels in patients. Given the reported reduction in oligodendrocyte density in schizophrenia(24), we expected the microstructural changes of increased qT1 and AxD in patients with untreated first episode psychosis. Furthermore, on the basis of Tandon’s hypothesis that higher cholinergic tone underlies greater negative symptoms(46), and the cholinergic hypothesis of mania(47), we tested if lower qT1 (higher myelin) of cholinergic nuclei could predict higher negative symptom burden and lower manic symptoms. Given previous links between psychiatric disorders and cholinergic dysfunction, we expected gene expression of the basal forebrain to show enrichment for schizophrenia-related genes.

Next, we tested covariance between qT1 of BF and cortical-subcortical regions for internal validation of the BF measures of microstructure, and externally validate the covariance maps using imaging-transcriptomics. We predict significant covariance between qT1 of BF and cortical-subcortical regions known to receive cholinergic projections and expect this spatial covariance to align with the expression patterns of cholinergic receptor genes. The basal forebrain cholinergic system is divided into magnocellular cell groups Ch1, 2, 3 and 4--Ch1 and 2 (medial septum and vertical diagonal band nucleus) are thought to project to the hippocampus, Ch3 to the olfactory bulb, and Ch4 to the cortex and amygdala(48, 49). For the hippocampus, we expect CA1-subiculum to demonstrate strongest correlations given it has a higher density of cholinergic neurons based on choline aceyltransferase (ChAT) expression(50), and both CA1 and subiculum receives projections from the BF(51). In the cortex, we expect strongest correlations within regions adjacent to the external capsule based on the neuroanatomy of the medial and lateral pathways from the Ch4(52). Tracer studies indicate frontal, entorhinal, and anterior cingulate cortices (53-55), and more recent work show medial prefrontal cortex, sensory and motor cortices, and entorhinal cortex as projection targets of the BF, and cortical cholinergic neurons were observed largely in layer 2-3 of the prefrontal, motor/somatosensory, and visual cortices(34)—therefore we expect frontal, entorhinal, and motor/sensory cortices to show significant covariance. In imaging-transcriptomic correlations of BF-cortical covariance, we expect enrichment for oligodendrocyte marker genes given qT1 measures myelin content and qT1-based correlations would be driven by myelin-related genes. We would also expect enrichment for cholinergic neurons given presence of cholinergic interneurons may also drive correlations between the cortex and BF. Amongst the genes, we expect significant associations with cholinergic receptor genes as regions with high correlations are more likely to receive cholinergic input from the BF.

## Materials and Methods

### Clinical participant recruitment and assessment

Details regarding recruitment have been described in our previous work (44, 56, 57) but included here for completion. Participants were recruited as part of a neuroimaging project tracking changes in early psychosis from the PEPP (Prevention and Early Intervention for Psychosis Program) at London Health Sciences Centre. All participants provided written, informed consent with approval from Western University Health Sciences Research Ethics Board. Inclusion criteria were as follows: individuals experiencing FEP, with lifetime antipsychotic treatment less than 14 days. Exclusion criteria for FEP included: meeting criteria for a mood disorder (bipolar or major depressive) with psychotic features, or possible drug-induced psychosis. Healthy control (HC) participants were free from personal history of mental illness or family history of psychotic disorders, matched based on age, sex, and parental education. Exclusion criteria for both FEP and HC included substance use disorder in the past year based on DSM-5 criteria, history of major head injury, significant medical illness, or contraindications to MRI. Participants were assessed using DSM-5 criteria and the 8 item Positive and Negative Syndrome Scale (PANSS-8), Young Mania Rating Scale (YMRS), the Calgary Depression Scale (CDS), and cannabis use evaluated using the Cannabis Abuse Screening Test (CAST).

### Acquisition of neuroimaging and preprocessing

Details regarding imaging acquisition parameters and preprocessing are provided in Supplementary Information—briefly, we acquired T1-weighted images (qT1), diffusion tensor imaging, and MRS measures of choline in the dorsal ACC. Preprocessing for T1-weighted images are in line with our previous work(57), as well as DTI(58) and MRS(56, 57). Following preprocessing, we used the T1-weighted images to map the cortex and subcortical structures using automated methods (see Supplementary Methods). The basal forebrain is defined using a published probabilistic atlas based on cytoarchitectonic mapping of cholinergic cell groups, Ch1-3 and Ch4 (NBM)(26), and warped to individual subject T1-weighted images using ANTs(59) (Supplementary Methods).

### Neuroimaging statistical analyses

In the case-control analysis, we compared mean qT1 of BF structures between HC and FEP, using multiple linear regression seeking the main effect of diagnosis accounting for age, sex, cannabis use (CAST), and smoking status as covariates. We tested for clinical correlates of the BF measures. Considering the cholinergic hyperactivity hypothesis of negative symptoms in schizophrenia(4, 6, 7, 46, 60), and the cholinergic-adrenergic hypothesis of depression and mania(47, 61), and given past evidence of ACHe inhibitors improving manic symptoms(62), we hypothesized that structural integrity of the BF, reflecting cholinergic tone, would be correlated with manic and negative symptom severity. We predicted that higher qT1 (low myelin) and thus reduced cholinergic tone would correlate with greater manic symptom severity, and that lower qT1 (high myelin) and increased cholinergic tone to predict greater negative symptom burden.

We correlated the BF microstructural measures with choline, expecting the BF to predict cholinergic tone. We also examined the interaction effect of diagnosis x microstructure, in that the relationship between structure and choline would be altered in FEP. In HC, we expected qT1 to correlate negatively with choline—such that lower qT1 (high myelin) would reflect higher cholinergic tone and elevated levels of choline.

### Neuroimaging correlations

We correlated qT1 values between the BF and the rest of the brain including the subcortical brain structures (hippocampus, amygdala, striatum, thalamus, globus pallidum) and cortex. This is based on previous work that shows cellular similarity, and connectivity as a basis for correlations between morphometric measures(63). We hypothesized that the most significant correlations would reflect 1) Brain regions receiving cholinergic input from the basal forebrain, and 2) Brain regions with relatively higher cholinergic neuron content. Pearson’s R correlations were corrected for multiple testing by FDR within each structure, with significance threshold of FDR q < 0.10.

### Transcriptomic analysis of basal forebrain

We examined gene expression profiles of the BF structures to determine whether transcriptomics would reveal enrichment for cholinergic cell type markers, and disease associations. We used gene expression data from the Allen Human Brain Atlas(64). Quality control was done by visual examination of BF samples on the MNI template. One Ch1-3 sample was found to be misplaced or mislabelled (see Supp Figure 1), and therefore excluded from analysis. For the NBM, 4 donors contributed 9 NBM samples and the same 4 donors contributed 8 Ch1-3 samples (nucleus of the diagonal band, 4 horizontal and 4 vertical division) after quality control.

Probe selection was based on published quality control metrics based on comparison between the Agilent microarray and “ground truth” RNA-Seq(65). We selected those probes that passed quality control and selected one probe per gene based on highest correlation to RNA-seq data. Probe-to-gene mappings were provided by the Allen Institute.

We identified genes that were specifically expressed in the Ch1-3 and NBM. Region-specific expression analysis, as in previous studies(66, 67), was done using R (version 4.0) and the limma package to detect genes specifically expressed in the BF. For this analysis, all samples are included, restricted to only those donors that sampled the Ch1-3 and NBM.

For each gene, linear models were fit to gene expression with terms for the donor and region of interest. Moderated t statistics and p-values were calculated using the empirical Bayes moderation method, and corrected for multiple comparisons using Benjamini-Hochberg false discovery rate (FDR) (68). Genes were ranked based on significance (-log10 of p-value) multiplied by the sign of the t-statistic, and we selected the most significantly enriched genes after FDR correction at q < 0.10, 0.5, and 0.1. These gene lists were submitted to the Cell-type Specific Expression Analysis (CSEA), which we expect to show enrichment for cholinergic neurons(69, 70). ToppGene (71) was used to test for disease enrichment (at q < 0.10), using the DisGeNet Curated database, exploring whether there was enrichment for schizophrenia-related genes.

### Imaging-transcriptomic validation

We sought to explain BF-cortical imaging correlations with gene expression. We used data from the AHBA for the left cortex only since only 2 of the 6 AHBA brains sampled the right hemisphere. Only cortical samples are retained (samples from non-cortical, i.e., subcortical regions are removed), prior to finding a 1-to-1 match between each AHBA cortical sample and CIVET-generated vertex (Supp Figure 2a).

For matching AHBA samples to vertices on the CIVET cortical template, we used the set of re-registered AHBA sample coordinates to the MNI ICBM 2009c (nonlinear symmetric) template (generated by Gabriel A. Devenyi; https://github.com/gdevenyi/AllenHumanGeneMNI). This improves anatomical concordance between the original anatomical sample and MRI using multispectral and non-linear registration, as the original AHBA-provided MNI coordinates were calculated using an affine registration only. The ICBM template is also processed through CIVET 2.1.1 to generate a cortical surface. We assigned each AHBA sample to the closest vertex based on Euclidean distance. Where multiple samples matched to the same vertex, the closest sample in terms of distance was retained. We removed 2 samples with distance greater than 10mm, which were in the right hemisphere based on visual inspection (Supp Fig 2b). We matched 1,236 AHBA samples to a unique vertex, with mean distance of 1.47 between sample and vertex (sd 0.89, range 0.084 to 6.78mm).

Pearson’s R correlations for the left cortex (as described above) were modelled using mixed effects models, with gene expression as the fixed and donor as random effect. This is repeated across ∼ 14,000 genes and resulting p-values are corrected using FDR. Genes passing FDR threshold (at 10%) were submitted to CSEA.

## Results

### MRS-structural correlations

We tested for correlations between structural measures and dACC choline. There was no significant difference in dACC choline between HC and FEP (t= 1.08, p= 0.284, DF=78) after controlling for age, sex, smoking status and cannabis use.

In healthy controls (N=29), we found significant correlations between dACC choline and left NBM qT1 (Pearson’s R=-0.517, p=4.08E-03, DF=27), marginally so for Ch1-3 (R=-0.35, p=0.062), but not the right NBM (R=-0.238, p=0.2127) (Figure 1a). After accounting for influence of covariates including age, sex, and cannabis use, on choline, correlations continued to be significant--for Ch1-3 (t=-2.62, p=0.015, DF=23), and left NBM (t=-4.34, p=2.4E-04, DF=23). These correlations were not significant in the FEP group (p=0.292, 0.874, 0.307 for left, right NBM and Ch1-3 respectively).

**Figure 1.**
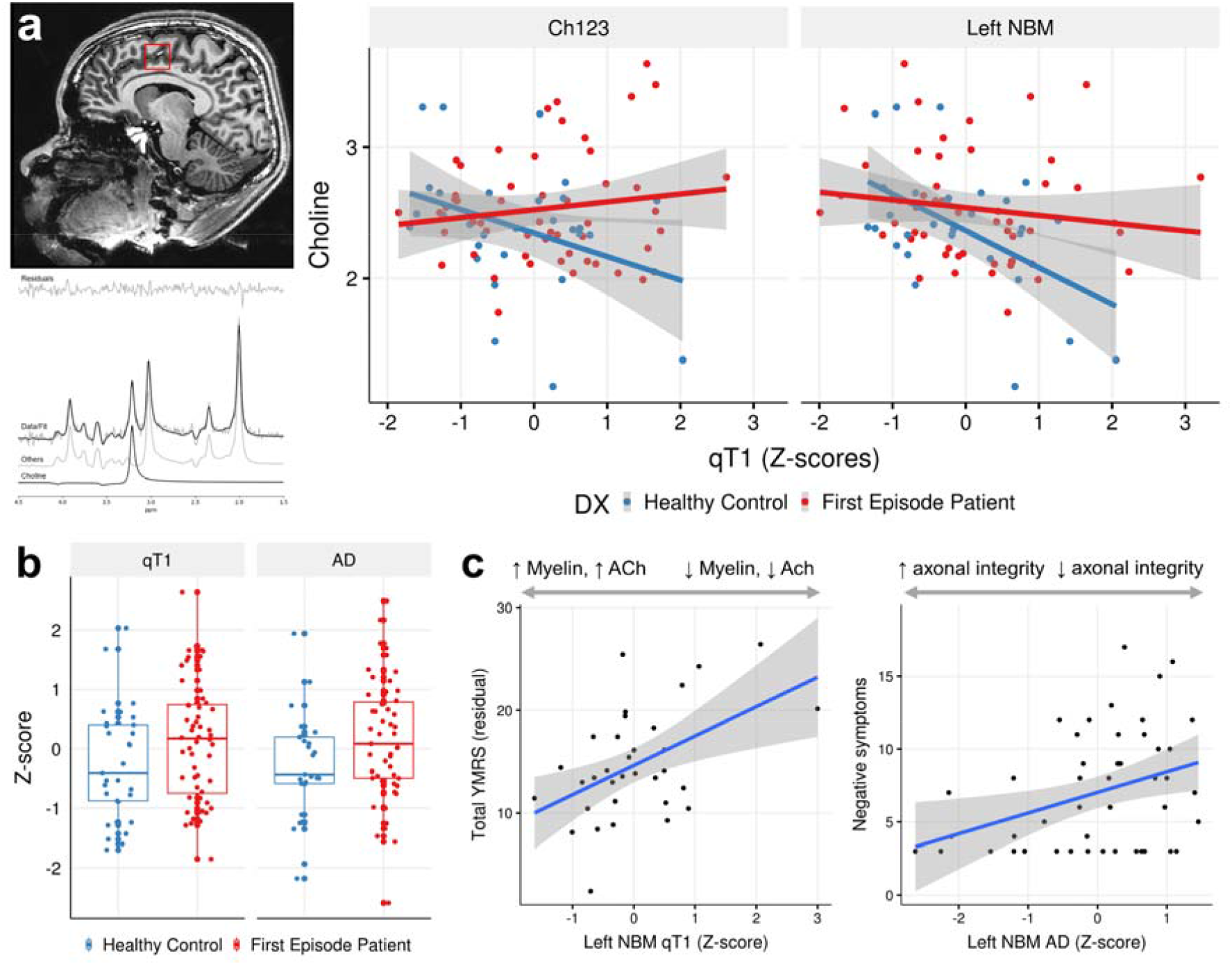
Evidence for microstructural changes of the basal forebrain in FEP. **a)** Left: Magnetic resonance spectroscopy of choline in the dACC with example of voxel placement (top), and spectral fit for choline (bottom). Right: In FEP, there is decoupling of the correlation between qT1 and choline levels, while a significant correlation exists in healthy controls such that lower qT1 (higher myelin) is associated with elevated choline levels. **b)** In the Ch1-3, there is increased qT1 (lower myelin), and increased axial diffusivity. c) Higher qT1 (indicating lower myelin) of the left NBM is associated with greater manic symptom severity, and increased AxD (reflecting lower axonal integrity) is associated with negative symptom severity. Left: y-axis shows residuals of the linear regression (YMRS ∼ cannabis use) added to the mean YMRS score.

We tested for the differences in slopes by modelling the diagnosis-by-qT1 interaction effect on choline levels which showed significant differences for Ch1-3 (t=2.96, p=4.14E-03, DF=76), and left NBM (t=2.72, p=8.10E-03, DF=76) after accounting for age, and sex as covariates (Figure 1a). Smoking status and cannabis use were excluded in interaction analysis since the healthy control group had no smokers, and the FEP group had significantly higher cannabis use (t=5.63, p=4.50E-07, DF=64).

### Case-control differences in BF microstructural measures

This analysis includes N= 85 (29 HC, 56 FEP), using multiple linear regression to examine the main effect of diagnosis on microstructural measures of the BF, including qT1 and AxD. We found significant differences between HC and FEP in Ch1-3 qT1 (t= 2.717, p=8.64E-03, DF=59) after accounting for age, sex, cannabis use, and smoking as covariates (Figure 1b). There were no differences in left or right NBM (p > 0.10). For the Ch1-3, mean AxD was increased in FEP (t= 2.90, p=5.58E-03, DF=50) with the same covariates (Figure 1b). These results for Ch1-3 persist after including ICV as a covariate--for qT1 (t=2.88, p=5.57E-03, DF=58) and AxD (t=2.94, p=5.02E-03, DF=49).

Overall, we found evidence for increased qT1 and AxD in the Ch1-3 (medial septum and diagonal band) of the basal forebrain in FEP, independent of brain size.

### Basal forebrain microstructure in relation to clinical scores

We sought clinical correlates of BF structural measures in the FEP group (N=56). We examined the left NBM and Ch1-3 along with measures of qT1 and AxD given the MRS findings above. We correlated structural measures with total YMRS and negative symptom scores.

There was a significant correlation between left NBM qT1 and YMRS (t= 3.03, p=5.09E-03, DF=29) after accounting for cannabis use and smoking status (Figure 1c), accounting for 24.7% (adjusted R-squared) of the YMRS variance. Ch1-3 axial diffusivity correlated with YMRS neared significance (R=-0.286, p=0.054), and less significant after accounting for cannabis use and smoking status (p=0.90). Left NBM AxD was correlated with negative symptom severity (R=0.358, p=0.0115, DF=47), remaining significant after accounting for smoking status and cannabis use (t=2062, p=0.0490, DF=27). After multiple testing correction (2 structures x 2 metrics x 2 clinical scores= 8 tests), only the left NBM qT1 to YMRS correlation is significant (p_Bonf_ < 6.25E-03).

### Neuroimaging correlations

Outlined in Figure 2 is the neuroimaging preprocessing, including cortical and subcortical segmentation and sampling (Figure 2a), registration of the probabilistic atlas to subject space (Figure 2b, top) and resulting distribution of mean qT1 values as an example (Figure 2b, bottom). Figure 2c shows range of Pearson’s R values and thresholded (after FDR correction) regions from BF-cortical and BF-hippocampal qT1 correlations. For BF-cortical correlations, sample size was N=65 (22 HC, 43 FEP) after quality control of CIVET cortical surfaces, and for BF-hippocampal correlations, N=76 (27 HC, 49 FEP) after QC for subcortical segmentations.

**Figure 2.**
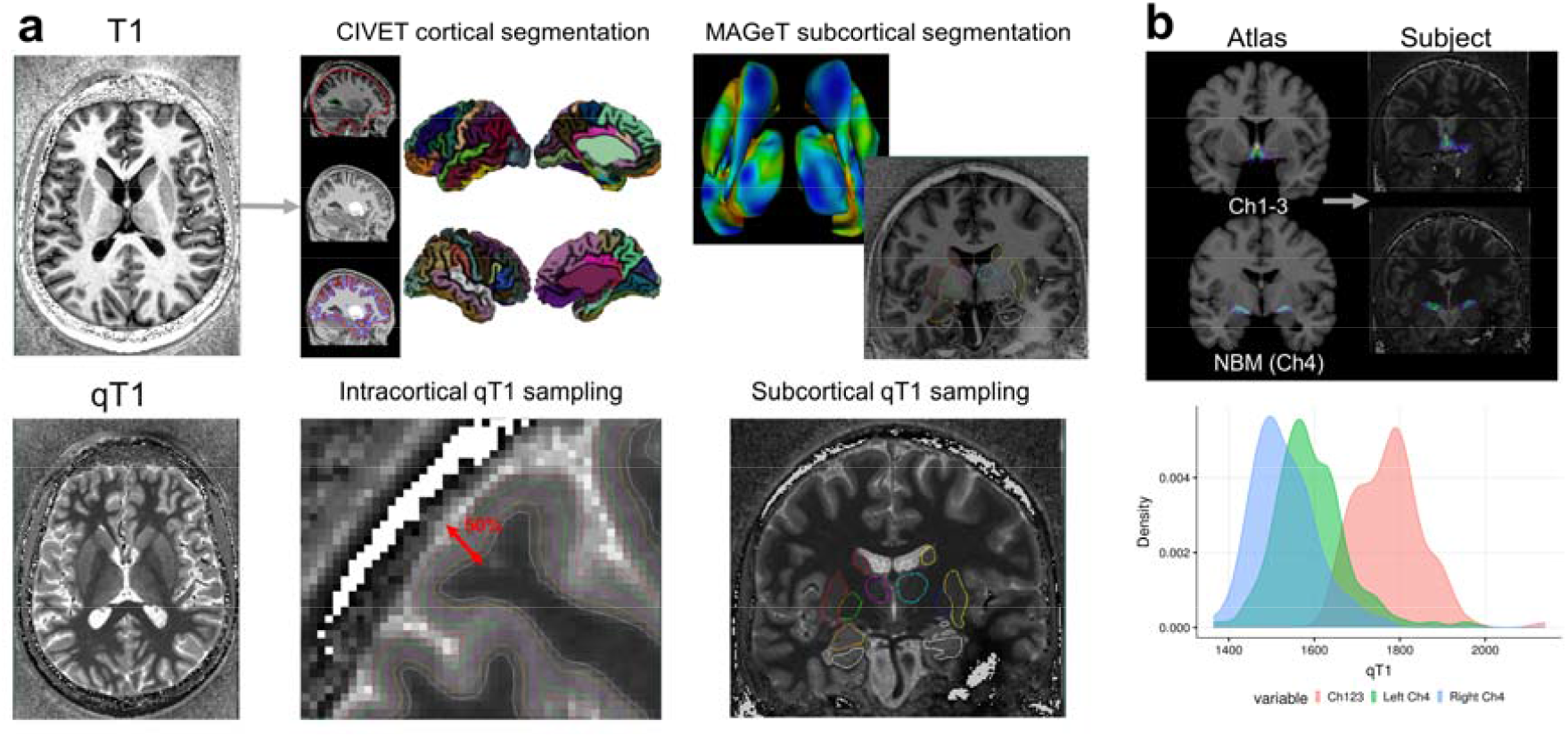
Image preprocessing of cortical, subcortical, and basal forebrain structures. **a)** qT1 and T1-weighted MRI data at 7T were acquired using the MP2RAGE sequence. The CIVET pipeline (version 2.1.0) was used to delineate the cortical surfaces and sampling at multiple depths, and we sample qT1 measures at 50% depth from the pial to white matter surface. Subcortical qT1 sampling along MAGeT Brain-generated structures (hippocampus, amygdala, striatum, thalamus, globus pallidum). **b)** Mapping probabilistic atlas based on cytoarchitectonic data of the basal forebrain structures onto individual subjects through non-linear registration. Histogram shows group distribution of qT1 values for the three structures—Ch1-3, left and right Ch4 (NBM).

Correlating qT1 values between the BF and cortex demonstrates the known projection pathways from the BF (Figure 3a). The medial cortical surfaces show significant correlations, reflecting the medial pathway of Ch4 projections(52). After FDR correction, we observed significant correlations for frontal (Figure 3a, blue arrow), precentral/postcentral (red arrow). BF-hippocampal correlations show the CA1 and subiculum as the most prominent regions (Figure 3b, green arrows).

**Figure 3.**
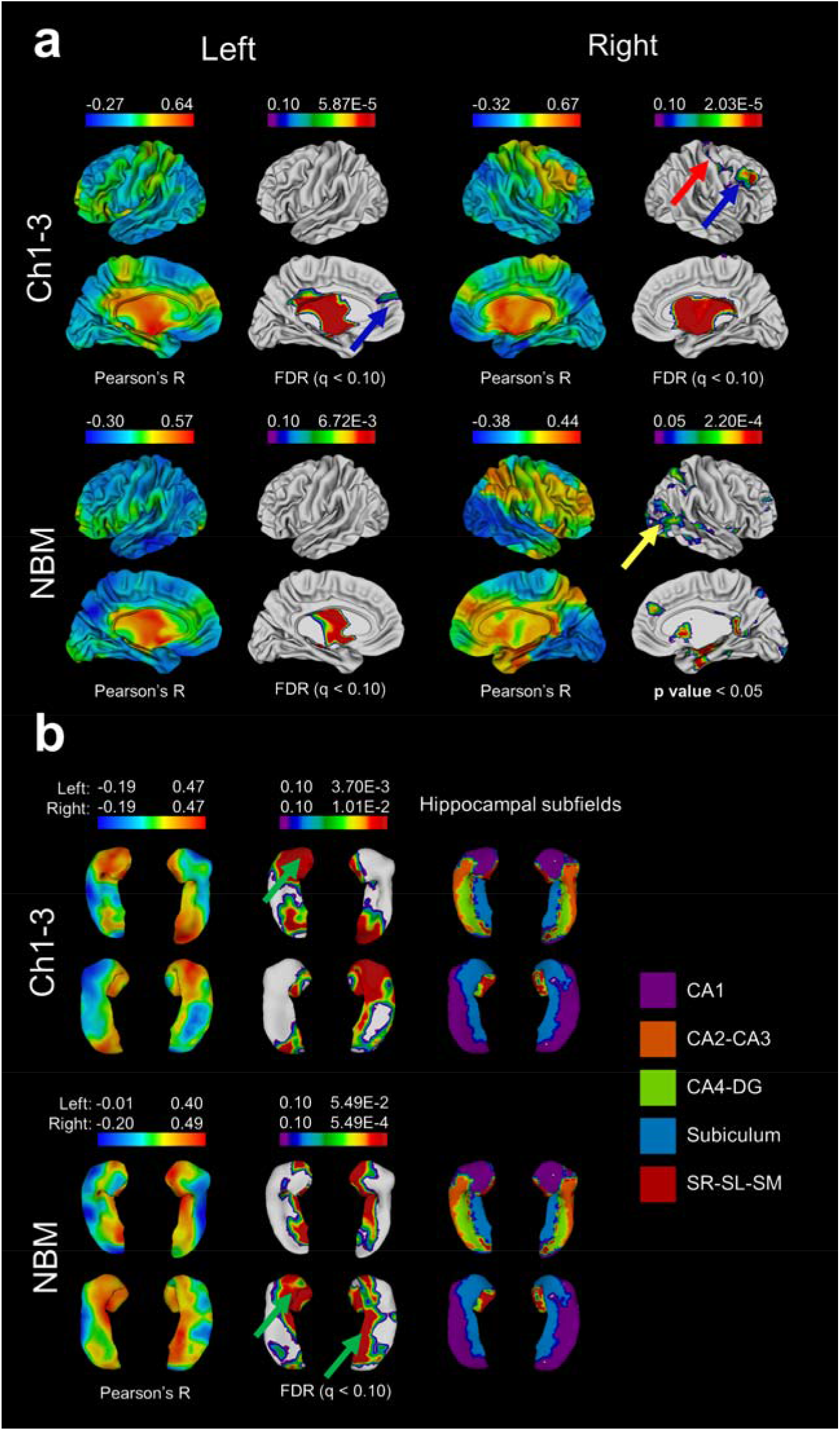
NBM-cortical and NBM-subcortical qT1 correlations for the a) Cortex and b) Hippocampus. Warm colours (red) indicate most positive correlations, and cool colours (blue) negative correlations for each set of surfaces. Regions surviving FDR are shown. Findings are consistent with the hypothesis with strongest (most positive) correlations reflecting cortical regions receiving the most consistent afferent projections the BF, and those regions with cortical cholinergic neurons. These include the frontal (blue arrow), sensory/motor cortices (yellow arrow). For the hippocampus, CA1-subiculum demonstrate strongest correlations (green arrow).

### Transcriptomic analysis of BF structures

We examined genes with specific expression in the Ch1-3 and NBM, at FDR corrected thresholds of 10, 5, and 1%.

ToppGene disease enrichment shows Ch1-3 (at FDR 10%) genes showing significant overlap with schizophrenia-related genes (1226 genes from Ch1-3, 79 genes overlapping with 883 schizophrenia-related genes) (q=4.744E-03) (Figure 4a). In the NBM, 2880 genes survived FDR 10%, and 153 genes overlapped (q=2.51E-02). For both sets of overlapping genes, we found 4 acetylcholine-related genes including *CHAT, CHRM2, ACHE*, and *CHRNA3. CHAT* was the most significantly expressed gene in both Ch1-3 and NBM out of the intersecting genes, and *ACHE* was amongst the top 10 genes. CSEA showed enrichment for cholinergic neuron markers in the Ch1-3 and NBM (Figure 4b).

**Figure 4.**
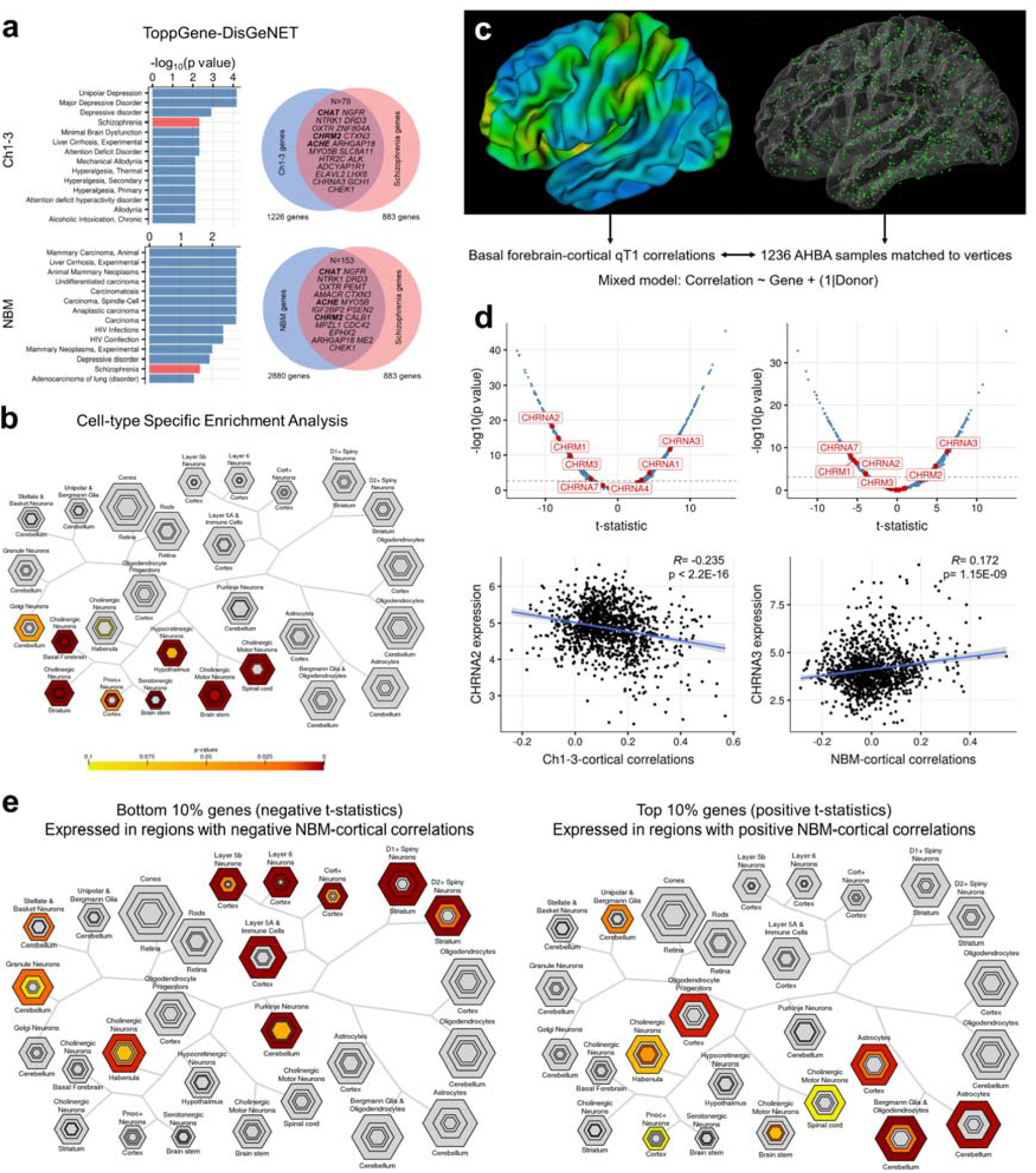
Transcriptomic analysis of the basal forebrain structures (a-b) and using transcriptomics to explain neuroimaging correlations (c-e). **a)** Enrichment of schizophrenia related genes. Venn diagram shows overlap between genes highly expressed in Ch1-3 and NBM with schizophrenia-related genes based on DisGeNET. Bolded genes are associated with cholinergic function (*CHAT, ACHE, CHRM2*). **b)** CSEA of genes passing FDR (q < 0.10) shows enrichment for cholinergic neurons in the Ch1-3. **c)** Coloured cortical surface (left) shows Pearson’s R correlations between Ch1-3 qT1 and intracortical qT1. On the right surface, 1236 AHBA cortical samples assigned to a unique vertex are shown. Using a mixed effects model, we sought to explain BF-cortical correlations by gene expression with donor as a random factor. **d)** Top: Volcano plot with distribution of t-statistics (x-axis), with dotted line indicating significance at FDR q < 0.05. Labelled genes are cholinergic receptor genes significant after FDR correction, for the Ch1-3 (left) and NBM (right). Bottom: example of imaging-genetic correlations for 2 of the most significant genes. **e)** CSEA of the top 10% (highly expressed in regions with positive NBM-cortical correlations) and bottom 10% genes (highly expressed in regions with negative NBM-cortical correlations) highlights relative enrichment of glial cells in regions with positive imaging correlations.

**Figure 5.**
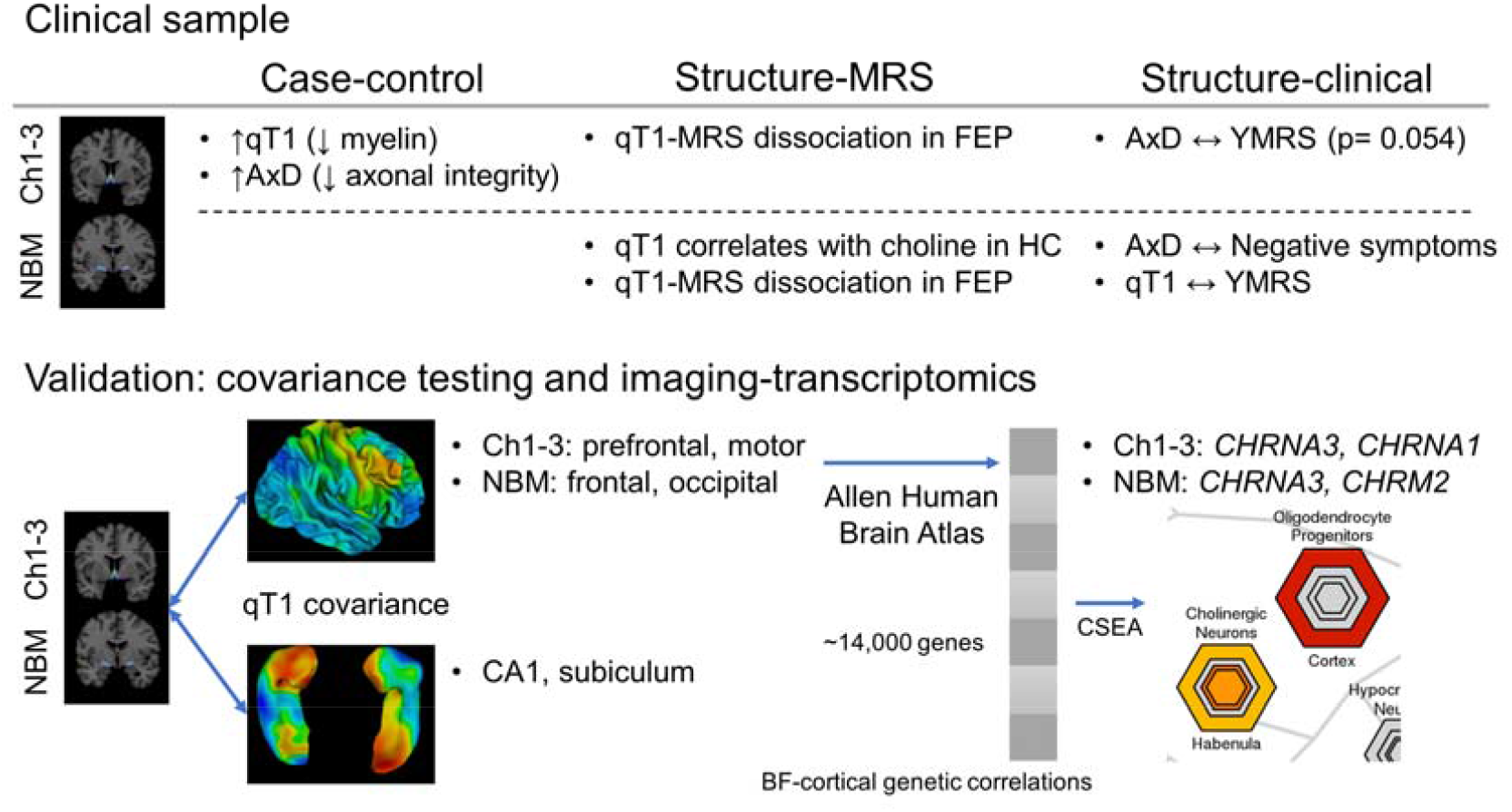
Summary of results, 1) Clinical sample neuroimaging results examining case-control differences in microstructure, structure-MRS relationships, and clinical correlates of microstructure. We found increased qT1 and AxD in FEP compared to healthy controls (HC). In HC, there was a significant correlation between qT1 (left NBM) and MRS choline, while left NBM was related to negative (AxD) and manic symptom (qT1) burden. 2) Validation of basal forebrain microstructure measures using qT1 covariance between cortical and subcortical regions, and further external validation of cortical covariance by correlating with gene expression using the Allen Human Brain Atlas. Genes outlined are cholinergic receptor genes most positively associated with BF covariance for Ch1-3 and NBM, surviving FDR correction.

### Imaging-transcriptomic analyses

We sought to explain BF-cortical correlations via gene expression. Using mixed effects analysis, we modelled the BF-cortical qT1 correlations by gene expression (per gene) and donor as a random effect (Figure 4c). At varying thresholds of FDR (10, 5, 1%), cortical qT1 covariance of NBM was significantly associated with 2262, 1513, and 754 genes respectively, and 4087, 3343, and 2278 genes for Ch1-3-covariance. We found a significant spatial correspondence between BF-cortex qT1 covariance and the distribution of cholinergic receptor gene expression (Figure 4d), with *CHRNA3* being positively correlated (i.e. highly expressed in those cortical regions with strongest correlations with Ch1-3 and NBM) (Figure 4d). *CHRNA2* was negatively correlated (i.e. highly expressed in cortical regions with negative correlations with BF) (Figure 4d). We used CSEA to interpret the imaging-genetic correlations by testing the top and bottom 10% percentile genes (most extreme t-statistics). For NBM-cortical correlations, CSEA of the top 10% (1512) genes (indicating positive t-statistics) shows enrichment for glial cells (astrocytes and oligodendrocytes) and cholinergic neurons, while the bottom 10% shows enrichment for cortical neurons (Figure 4e). Ch1-3-cortical correlations showed similar results (Supplementary Figure 3).

## Discussion

Studying the microstructure of the BF cholinergic nuclei for the first time in first-episode psychosis (FEP), we report an increased qT1 (reflecting reduced myelin content) and increased AxD (reflecting an increase in non-myelinated cellular fraction, and reduced axonal integrity) of anteromedial nuclei (Ch1-3) in patients. While dorsal ACC choline levels reflected the microstructure of NBM in healthy individuals, this relationship was absent in patients, indicating a possible dissociation between the cholinergic inputs to the Salience Network in psychosis. Our findings agree with the cholinergic-adrenergic hypothesis of mania, and possibly Tandon’s hypothesis that higher cholinergic tone underlies greater negative symptom burden. The transcriptomic analysis of the cholinergic nuclei further supports these findings by highlighting cholinergic neurons and dysfunction in schizophrenia and depression.

While illness-related effects were most pronounced in the anteromedial BF (Ch1-3), the variance in NBM microstructure related to cortical MRS choline, and clinical severity of manic and negative symptoms. This may indicate that a diffuse cholinergic deficit involving hippocampal projections may underlie psychosis, but more extensive deficit involving cortical projections may worsen the symptom burden. This distinction is based only on correlation analyses; studies with longitudinal design are needed to test this speculation. In testing structure-MRS correlations, we found left NBM had stronger correlations to dACC choline than the Ch1-3, which may be due to the NBM containing the highest proportion of cholinergic neurons (at least 90%) compared to Ch1 (10%), Ch2 (70%), Ch3 (1%) in the rhesus monkey (72). Further, left NBM qT1 and AxD were correlated with manic and negative symptom burden, suggesting the left NBM being most predictive cholinergic function across healthy controls and FEP.

We used imaging correlations as a basis for validating qT1 as a reliable measure for basal forebrain microstructure. We found significant qT1-based correlations between the BF cholinergic nuclei and cortical (frontal, precentral/postcentral), and subcortical regions (CA1 and subiculum in the hippocampus). Regions with strongest correlations seemingly reflected the known projection patterns from the BF and presence of cholinergic neurons, in particular the hippocampus with CA1-subiculum (50, 51). We found significant cortical correlations along the expected medial and lateral pathways from the Ch4(52), or along the medial cortical surfaces (Figure 2c). Significant cortical regions including frontal, sensory/motor, and visual cortices are highlighted in previous tracer studies(34, 53-55). Further biological validation for the imaging correlations are provided by significant associations with cholinergic receptor genes (Figure 3d) and enrichment for both oligodendrocytes and cholinergic neurons (Figure 3e). The contrast of the top vs. bottom percentile genes highlights positive enrichment for glial cells, or in other words cortical regions positively correlated with the basal forebrain may house greater density of oligodendrocytes, and overall myelin correlations could be reflective of underlying glia-neuron ratios. Taken together, these analyses strengthen the validity of microstructural measures (qT1, AxD).

Despite the high field strength (7T), in our structural neuroimaging protocol we found very little grey-white matter contrast in the BF region therefore limiting confidence in automated techniques for grey-white matter classification such as VBM, and even less so in measuring its differences across individuals. For example, a recent study with similar methods (using the same cytoarchitectonic atlas) found reduced gray matter integrity, while findings were not significant after accounting for global gray matter volumes(16)—suggesting findings were not region-specific and ascribed to global changes. Here, in exploration of an alternative approach and using untreated samples, we demonstrated feasibility and biological validity of microstructural measures using both imaging-imaging and imaging-transcriptomic correlations, and results were not impacted by global effects. Our results are further supported by the previous finding of myelin maps as superior in representing cortical circuitry and gene expression compared to other metrics such as cortical thickness(73).

Analysis of basal forebrain gene expression identifies enrichment of schizophrenia genes in both Ch1-3 and NBM, and highlights cholinergic neuron markers (*ACHE, CHAT*). This highlights an understudied component of schizophrenia pathophysiology, and an update building on previous hypotheses and findings(24, 74). Our results are in line with neuropathological studies, yet there are limited past transcriptomic studies of the BF cholinergic nuclei. While not a focus of the study, we found enrichment of genes associated with major depressive disorder within the BF. The evidence base for cholinergic system involvement is stronger for depression than schizophrenia—for example, a PubMed search for (cholinergic) AND (depression) shows 9,871 results while (cholinergic) AND (schizophrenia) yields 3,205 results (as of May 2^nd^, 2021). As mentioned above, the cholinergic-adrenergic hypothesis of depression (47) along with recent clinical studies has led to renewed interest of the cholinergic system in psychiatric disorders(75, 76). The biological overlap of cholinergic dysfunction across depression and schizophrenia(77) signals a common dimension that cuts across diagnoses that may explain clinical overlap (such as negative symptoms in schizophrenia that mimic depressive symptoms)—indicating a need to refine our understanding of cholinergic networks to better guide treatment options in psychiatry.

Our study has a number of strengths as well as limitations. We studied the cholinergic profile of patients experiencing first episode psychosis (mean duration of illness = 31.13 weeks), with minimal or no antipsychotic exposure (lifetime antipsychotic exposure of mean 1.32 DDD equivalents, amounting to < 2 days of exposure to minimal effective doses of antipsychotics) and no exposure to anticholinergic drugs. We quantified nicotine use and adjusted for the observed variance in our analysis. Nevertheless, we could not quantify choline resonance from the BF due to technical limitations of MRS from this anatomical area. Furthermore, we were limited in the number of female participants in this study; we urge readers to exercise caution when generalising our results to female participants.

## Acknowledgements

Computations were performed on the Niagara supercomputer at the SciNet HPC Consortium (Loken et al., 2010). SciNet is funded by: the Canada Foundation for Innovation; the Government of Ontario; Ontario Research Fund -Research Excellence; and the University of Toronto.

We thank Dr. Joe Gati, for their assistance in data acquisition and archiving. We thank Dr. William Pavlovsky for consultations on clinical radiological queries. We thank Drs. Raj Harricharan, Julie Richard, Priya Subramanian and Hooman Ganjavi and all staff members of the PEPP London team for their assistance in patient recruitment and supporting clinical care. We gratefully acknowledge the participants and their family members for their contributions. The MRS pulse sequence package used was developed from a source code provided by Uzay E. Emir, PhD, and Dinesh K. Deelchand, PhD, Center for Magnetic Resonance Research (CMRR), Minneapolis, MN. It is based on original work by Gülin Öz and Ivan Tkáč. Our special thanks to Uzay E. Emir, PhD, Dinesh K. Deelchand, PhD, Gülin Öz, PhD, Ivan Tkáč, PhD, and Edward Auerbach, PhD, for their help with sequence testing and for valuable technical discussions and to Dennis W. J. Klomp, PhD, and Vincent O. Boer, PhD, for providing the asymmetric excitation pulse used in the 7T version of the sequence. We also acknowledge CFREF BrainsCAN funding that we received providing a reduced hourly rate for the MRI scans.

